# Fitness advantage of sequential metabolic strategies emerges from community interactions in strongly fluctuating environments

**DOI:** 10.1101/2024.06.14.599039

**Authors:** Zihan Wang, Yu Fu, Akshit Goyal, Sergei Maslov

## Abstract

Microbes growing in fluctuating environments employ two key metabolic strategies: sequential (diauxic) utilization and co-utilization of nutrients. Most work has focused on understanding and comparing these strategies physiologically for the growth of single species, rather than ecologically for the assembly of complex natural communities. This is in part because of the lack of good metrics quantifying the fitness of different metabolic strategies in ecological contexts. Here, we present a new consumer-resource framework that incorporates dynamic proteome reallocation, and use it to compare the fitness of metabolic strategies during community assembly. We introduce two notions of fitness of a strategy in fluctuating environments: the time-averaged growth rate and the biomass-weighted prevalence of microbes using a given strategy. We find that sequential utilizers, although disadvantaged in pairwise competitions, gain a significant edge during community assembly — an advantage that becomes more pronounced with increasing community diversity and the size of the species pool from which they are assembled. Low diversity communities resemble pairwise competitions and are dominated by co-utilizers, whereas high diversity, mature communities are dominated by the sequential utilizers. This shift is driven by two factors: the difference in lag times and the increased structural stability conferred by sequential strategies. Overall, our work provides several testable predictions about the co-occurrence patterns of microbes using different metabolic strategies.

## Introduction

Microorganisms use a variety of metabolic strategies to utilize resources in their environments. They typically either consume the available resources all at once (co-utilization) or one after another (sequential utilization) [1, 2, 3]. Both strategies have different effects on physiology, i.e., on the growth rate of a single species in isolation. Even in this context, the answer to which strategy is superior remains disputed. Sequential strategies have been theoretically shown to be growth-optimal for certain pairs of resources and co-utilization optimal for others, which depends on the position of the resources in the central metabolic network [4]. However, experimental studies [5, 6, 7, 8, 9] show that microbes often deviate from growth-optimality, by allocating part of their proteome to resources that are not currently utilized [10].

While understanding the advantages of using different metabolic strategies when species are grown in isolation is important, microbes in natural environments rarely grow alone. Instead, they live in complex multi-species ecosystems, where emergent factors that are distinct from growth rate alone may favor some strategies over others. These ecological effects have never been systematically studied, either experimentally or theoretically. Thus, studies of both strategies in an ecological context may shed light on how and why they may coexist in natural environments.

An intuitive understanding of how ecological contexts might change the competitive advantage of a metabolic strategy is as follows. When growing in isolation, species can always count on resources being present. In contrast, in communities, some resources may be depleted by other species before a species gets a chance to consume them [11]. This becomes only more likely as the number of species (community diversity) increases. Such an effect may lead to differences in the performance of a strategy between pairwise competitions and community contexts.

Fluctuating environments such as in serial dilution experiments — where resource availability strongly changes with time — present a natural scenario in which to compare sequential and co-utilizing strategies. Indeed, in steady-state environments such as chemostats, resources reach low concentrations, where even sequential utilizers become co-utilizers [1, 3].

Here, using simulations and theory, we quantify the advantages of metabolic strategies in complex communities using two definitions of fitness in fluctuating environments: namely the time-averaged growth rate and the biomass-weighted prevalence of microbes using a given strategy. We find that sequential strategies have a systematic fitness advantage over co-utilizing strategies, which emerges in community contexts, getting stronger with increasing community complexity (number of species) and the size of the species pool from which communities are assembled. In our simulations, low diversity communities are dominated by co-utilizers, while high diversity communities are dominated by sequential species. We then investigate the source of this behavior, and find two key contributors. They are (1) the decreased importance of lag time differences between sequential utilizers and co-utilizers in ecological contexts, and (2) increased resilience to resource fluctuations (structural stability [12, 13, 14]) of communities enriched in sequential utilizers. Our results have implications for how both sequential and co-utilization strategies coexist in nature.

## Results

### Incorporating proteome allocation to study metabolic strategies

There are two major classes of metabolic strategies: (1) co-utilizing strategies where species consume multiple available resources simultaneously, and (2) sequential or diauxic strategies where species consume resources one at a time according to a species-specific hierarchy. The impact of utilizing either of these strategies on a species’ growth rate has been extensively studied in the context of microbial physiology [4, 10]. One extreme point of view suggests that microbes optimize their growth rates on the currently available resources without allocating any proteome towards enzymes necessary to consume future resources [4]. This point of view leaves species vulnerable to long lag periods of no growth when resource availability changes, as may frequently occur during boom and bust cycles in nature and serial dilution experiments [5, 15]. The other point of view [10] cautions against pure optimality arguments using evidence showing that “a large fraction of proteomes is allocated towards proteins that are unnecessary for growth” including enzymes and transporters required to utilize resources that are not currently consumed. Indeed, experimental evidence in *E. coli* co-utilizing certain pairs of resources suggests similar allocation among enzymes used for each of these resources regardless of their current availability [16]. To capture the spectrum of possibilities with these two being extremes [4, 16], here we introduce a pre-allocation parameter *ϕ*_pre_ quantifying the fraction of the proteome always allocated towards the utilization of any resource regardless of their presence in the growth medium (Fig. 1c). Furthermore, in agreement with previous work, we account for dynamic allocation between the ribosomal and metabolic sectors of the proteome [16] (Methods). Each species is characterized by a set of randomly-selected growth rates in single resource environments, which are subsequently used to infer their growth rates in mixed resource environments (Methods). Importantly, we assume that growth rates in different single resource environments are selected independently (Methods). Thus, there is no trade-off between species’ growth rates in different resource environments.

**Figure 1:**
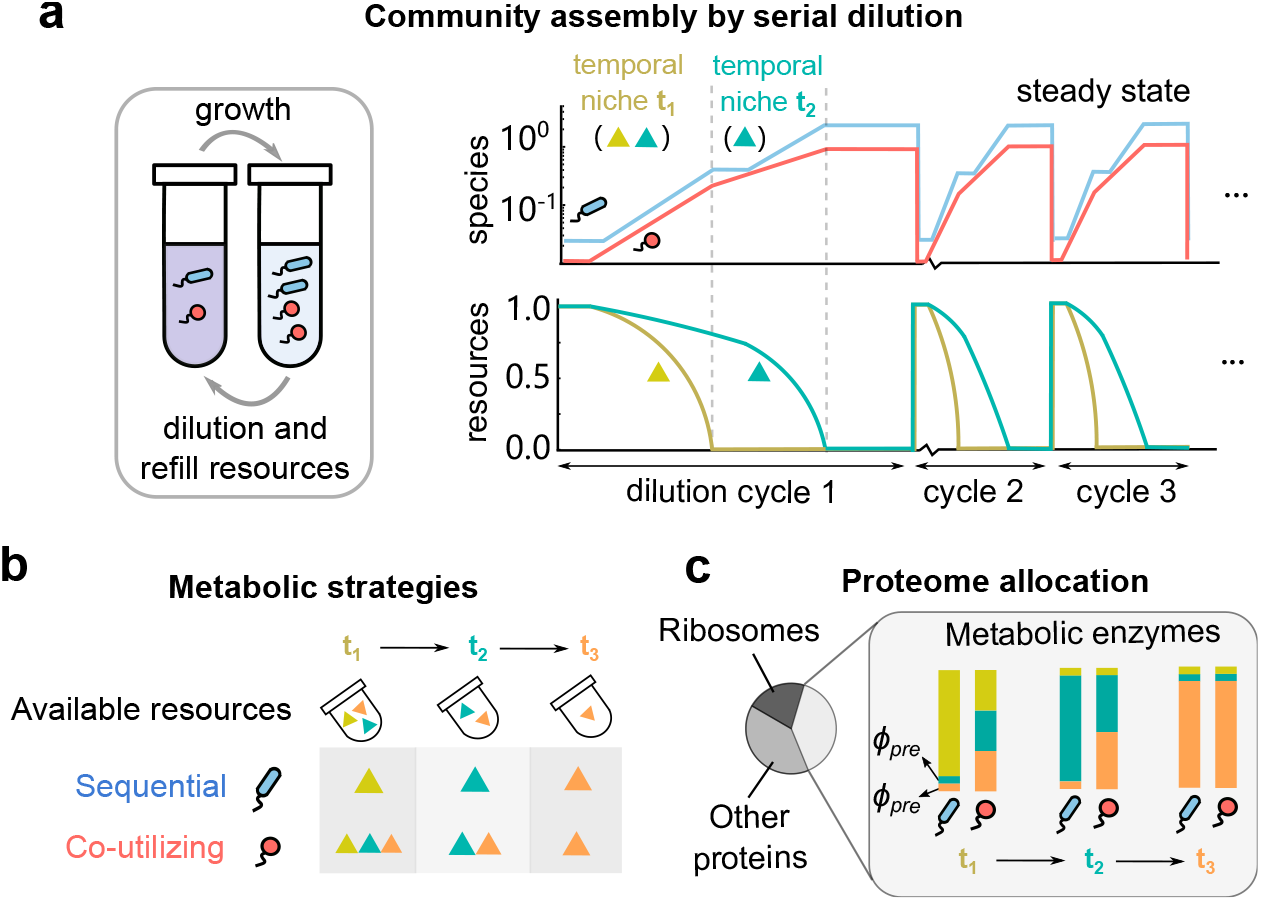
Investigating sequential and co-utilizing metabolic strategies in boom and bust environments requires incorporating proteome allocation in consumer-resource models. (a) Schematic illustration of species and resource dynamics in boom-and-bust (serial dilution) environments. In each growth-dilution cycle, resources are depleted in a specific temporal order, giving rise to “temporal niches” defined by which resources are currently available. Here, in the first niche lasting time *t*_1_, both resources (gold and teal) are present, while in the second niche lasting time *t*_2_, only one resource (teal) is present. After several growth-dilution cycles, both species (red and blue) reach a steady state where they grow by exactly the dilution factor *D* in each cycle. After a resource is depleted, species using it experience a lag time depending on how they preallocate internal enzymes (Methods). (b) Illustration of two major metabolic strategies: sequential (blue) and co-utilizing (red) used by the two species. Species using sequential strategies always consume one resource at a time, according to their idiosyncratically hard-wired preference order (here, gold, then teal, then orange). Co-utilizing species use all available resources that they can consume at the same time. (c) Illustration of proteome (re)-allocation in our model. We assume that each species has a proteome broadly divided into three sectors: ribosomes, metabolic enzymes and other housekeeping proteins. Within the metabolic sector, all species allocate a fixed small fraction *ϕ*_pre_ to each resource they are not currently utilizing. The rest of the metabolic sector is allocated according to the metabolic strategy: for sequential species all of the remaining metabolic sector is allocated to a single resource, while for co-utilizing species it is equally divided between all currently consumable resources.

One of the important roles played by pre-allocation is that it controls the lag times for switching between resource environments. A larger pre-allocation *ϕ*_pre_ typically shortens lag times, allowing species to quickly create the enzyme pools necessary for growth in a new environment. However, this comes at a cost of reducing the growth rate in the current environment since additional enzyme budget is used for pre-allocation [5, 9, 7, 6, 8]. In our model, we include this trade-off between growth rates and lag times (Methods) in accordance with experimentally verified models [5].

### Emergence of ecological advantage of sequential strategies

One way to compare different metabolic strategies is to measure the overall, or time-averaged growth rate of species utilizing them. This is straightforward when species are growing alone and can be done simply by taking the factor by which a species grows over a single boom-bust cycle (at steady state, this is always the dilution factor *D*) and dividing it by the time it takes deplete all resources (Fig. 2a). This metric closely resembles the notion of average fitness of a species, since it reflects a time-averaged growth rate. We measured the time-averaged growth rate for single species utilizing two specific strategies: co-utilization and the best sequential strategy identified in previous work [11] — the “top smart” one, where a species first utilizes the resource with the fastest growth rate, then uses all other resources in an idiosyncratic order uncorrelated with growth rate. We found that in single species contexts, co-utilizers have significantly higher time-averaged growth rates than sequential species (Fig. 2c), over the entire range of the pre-allocation fraction *ϕ*_pre_ tested. This observation can be attributed to the much lower lag times of co-utilizers compared with sequential species. Indeed the difference in time-averaged growth rates between strategies almost disappears if we unrealistically eliminate the lag times of both strategies (Fig. S5). This suggests that in single-species contexts, co-utilizers are more fit than sequential species.

**Figure 2:**
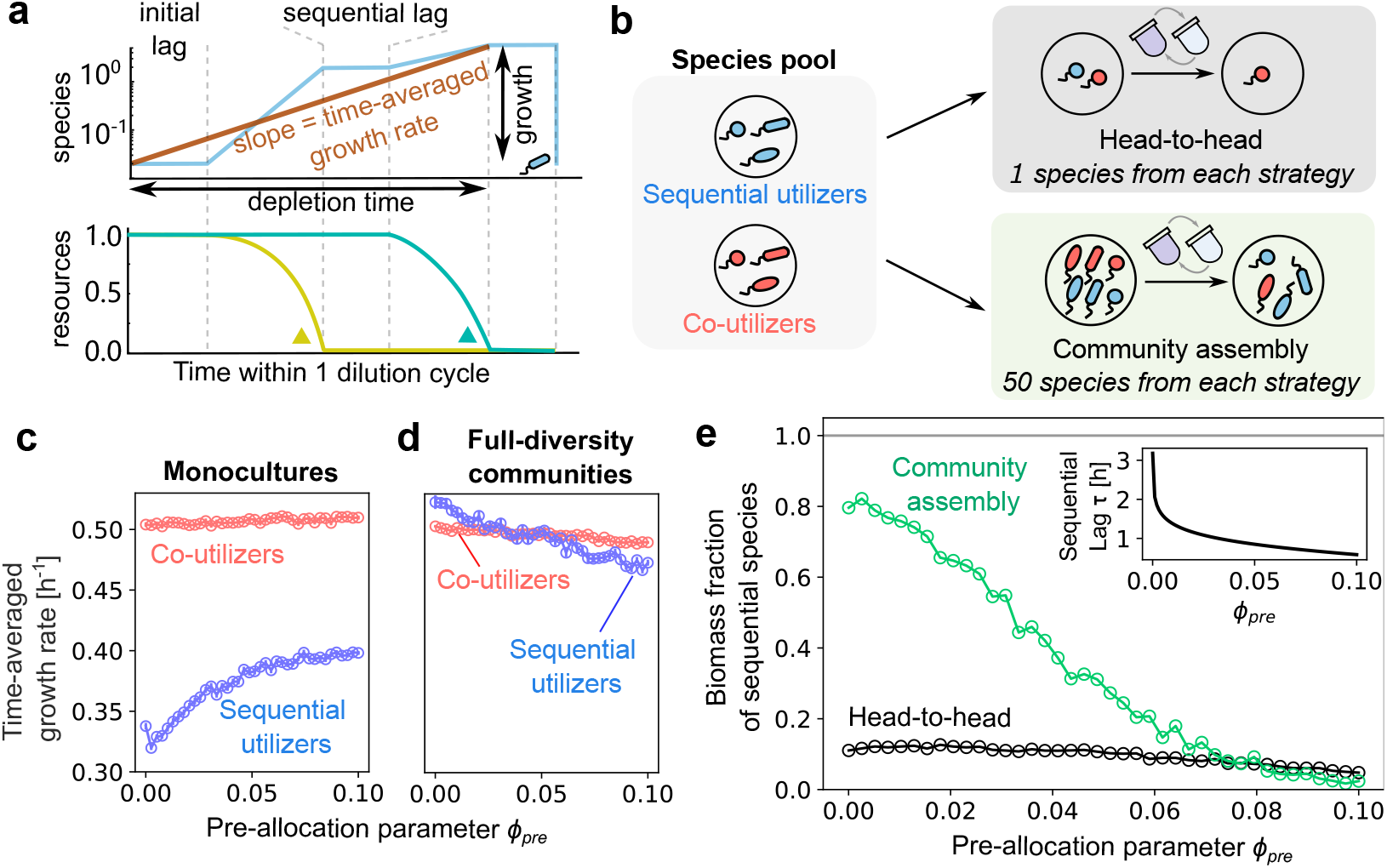
Comparison of co-utilization and sequential strategies in single-species and community contexts. (a) Schematic of time-averaged growth rate measurement. At steady state, the time-averaged growth rate is inversely proportional to the time it takes to deplete all resources. Top: species growth dynamics, including initial and sequential lags. Bottom: corresponding resource depletion over time. (b) Schematic showing our simulations in two different scenarios: one where we compete 1 sequential and 1 co-utilizing species head-to-head (gray), and the other where we assemble a complex community seeded with 50 species from each strategy (green). (c)-(d) Time-averaged growth rates of co-utilizers and sequential utilizers in monoculture and in “pure” communities of the same strategy with maximum diversity as a function of pre-allocation fraction *ϕ*_pre_. In monoculture (c), coutilizers consistently exhibit higher growth rates due to lower lag times. In full-diversity communities (d), sequential utilizers’ time-averaged growth rates become comparable to those of co-utilizers, and even exceed the latter at low *ϕ*_pre_. (e) Fraction of surviving sequential utilizers in each of the two scenarios: head-to-head (black) and in complex communities (green), plotted as a function of the pre-allocation fraction *ϕ*_pre_ common to all species in the pool. Sequential utilizers have a distinct ecological effect in complex communities: the green curve is typically above the black curve. Inset: average lag time *τ* during the first resource switch of sequential utilizers, as a function of *ϕ*_pre_.

However, in a community context, all surviving species must have the same time-averaged growth rate in order to coexist. Thus if co-utilizers and sequential species coexist, it will be impossible to compare them on the basis of their time-averaged growth rates. To overcome this challenge, we assembled “pure” communities comprising species utilizing only of the two strategies. To maximally distinguish community contexts from single-species contexts, we assembled communities of maximal diversity, where the number of species equaled the number of resources. Surprisingly, in such community contexts, the time-averaged growth rates of communities comprising co-utilizers and sequential species were comparable (Fig. 2d). At low values of *ϕ*_pre_, sequential species had a slight advantage in time-averaged growth rate, while at high values of *ϕ*_pre_, co-utilizers grew faster on average. This drastic reduction in the growth rate disparity between both strategies in ecological contexts can be explained as follows. As communities become more diverse, coexisting sequential species tend to be complementary to each other, i.e., they prefer different resources as their top choice [11]. Because of this complementarity, once species deplete their top choice resource, they often have no resources left to switch to since they have been depleted by others. Hence, the lag disadvantage of sequential species becomes essentially irrelevant in diverse community contexts. This allows them to become comparable in their time-averaged growth rates with co-utilizers. Hence, in community contexts sequential species can often be as fit, if not fitter, as co-utilizers.

Another way of comparing metabolic strategies is in terms of the prevalence of species utilizing them in naturally assembled communities. To this end, we assembled several multi-species communities, each starting from a large species pool with 50 species utilizing a sequential strategy and 50 species which were co-utilizers (Methods; Fig. 2b). We serially diluted these communities until they reached steady state. We measured the prevalence of sequential species as the biomass-weighted fraction of them among all survivors in assembled communities. We found that sequential species dominated in their prevalence (reaching as high as 85%) at low values of *ϕ*_pre_ — their prevalence gradually decreasing with increasing *ϕ*_pre_, reaching parity (50%) at *ϕ*_pre_ ≈ 0.03 (Fig. 2e). This confirms and amplifies the ecological effect we observed in time-averaged growth rates in Figs. 2c–d. Even the *ϕ*_pre_ values at which both strategies reach parity are comparable. Note that in contrast with the scenario in Fig. 2d, which was somewhat artificially constructed, the advantage of sequential species in Fig. 2f emerges naturally through the process of community assembly. This advantage persists in the absence of lag times, albeit it gets weaker and can be explained in terms of the statistics of growth rate distributions (Fig. S5). Together, these results point to the emergence of a distinct and significant advantage for sequential species in ecological contexts of species-rich communities.

### Sequential strategies promote diversity through increased structural stability

To better understand the forces guiding the assembly of complex communities of species using different metabolic strategies, we investigated the composition of communities with varying levels of species diversity or complexity. Because of competitive exclusion, the final species diversity ranged between 1 and the number of resources *n*_*R*_ = 4, depending on the randomly generated species pool. Interestingly, the fraction of sequential species was strongly correlated with this final species diversity (Fig. 3a). That is, across all values of *ϕ*_pre_ tested, sequential species dominated in high-diversity assembled communities, while co-utilizers were prevalent in low-diversity communities (Fig. 3a). Fig. 3b shows a detailed breakdown of the community diversity at different values of *ϕ*_pre_. As *ϕ*_pre_ increases, the most common community diversity shifts from maximally diverse (4 surviving species on 4 resources) to minimally diverse (1 surviving species). Further, at each value of *ϕ*_pre_, strategies were segregated by community diversity. That is, single-species communities almost entirely contained co-utilizers (≈ 100%) while maximally diverse communities with 4 species were dominated by sequential species (Fig. S6). This segregation of strategies by diversity was observed even at when *ϕ*_pre_ ≈ 0.036 (Fig. 3d), when sequential and co-utilizing species were on average equally prevalent in communities (Fig. 2f).

**Figure 3:**
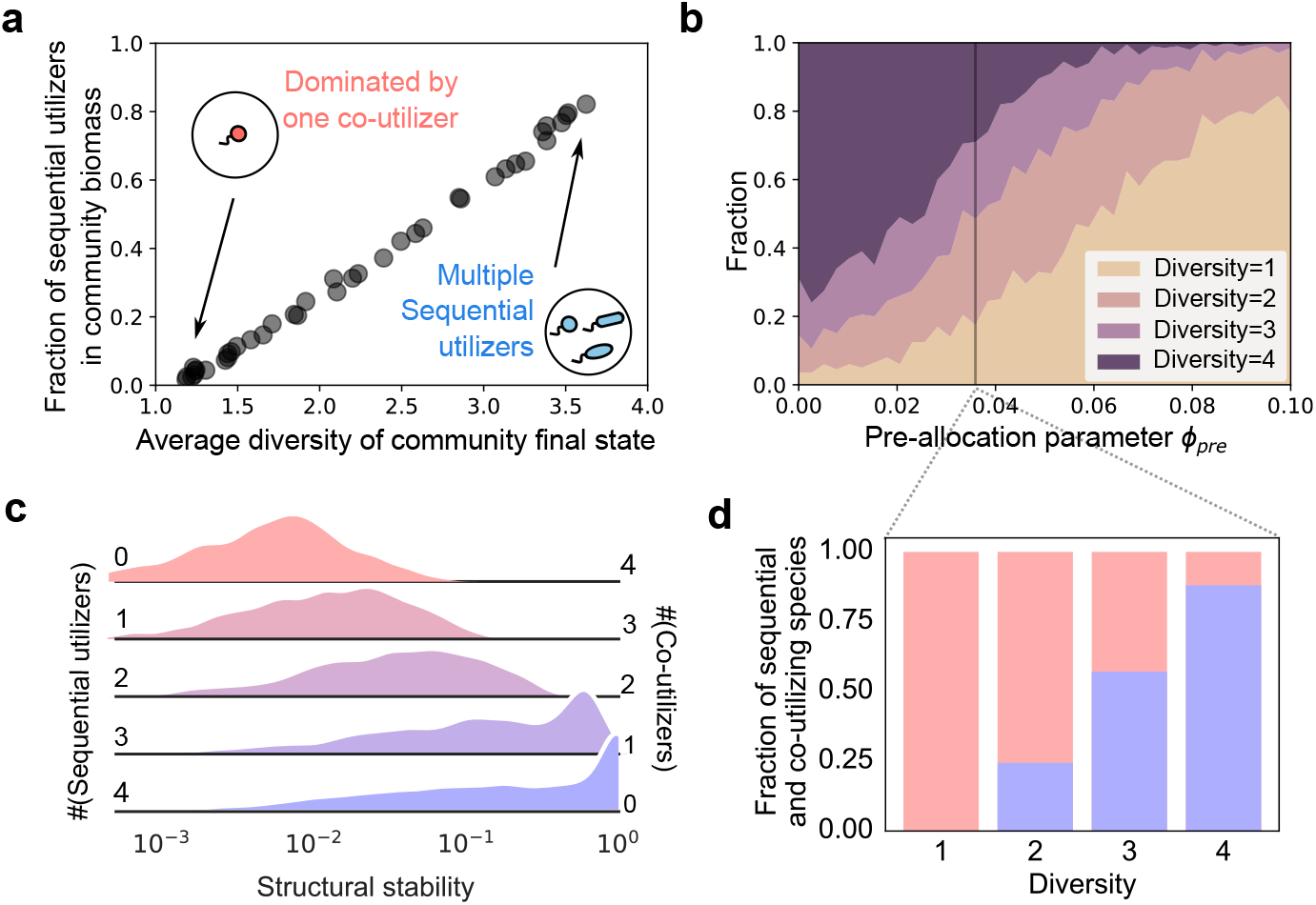
Sequential strategies promote diversity through increased structural stability. (a) Fraction of surviving sequential species in biomass as a function of the average diversity of the assembled communities (# of surviving species) for simulated communities across all *ϕ*_pre_ values tested. As community complexity increases, sequential species become more dominant in communities. (b) Stackplot showing how the diversity of assembled communities are stratified across *ϕ*_pre_. At low *ϕ*_pre_ the majority of communities have full diversity, while at high *ϕ*_pre_ most of them are dominated by 1 species. (c) Distributions of normalized structural stability (Methods) for niche-packed communities (*n*_*S*_ = *n*_*R*_ species). Increasing the number of sequential utilizers, from 0 to 4 systematically increases structural stability. (d) Stacked bar plot showing how sequential (red) and co-utilizing species (blue) are stratified across communities with different diversities at a fixed value of *ϕ*_pre_ ≈ 0.036. Here too, more complex communities are increasingly dominated by sequential species, while less complex communities are dominated by co-utilizing ones.

A plausible explanation for the stratification of strategies by community richness/diversity (number of species) lies in the systematic difference in their structural stability, i.e., the likelihood that a given community can successfully assemble in a given resource environment, e.g., when all resources are supplied in equal concentrations, as in our simulations. A given set of species may only stably coexist in a certain range of environmental conditions quantified by the supply of different resources (Methods). This range quantifies the structural stability [12], and we developed a linear algebra approach to calculate it for any niche-packed community (Methods). By quantifying the structural stability of niche-packed communities composed of different number of sequential species (from 0 to *n*_*R*_ = 4 at *ϕ*_pre_ ≈ 0.036), we found that structural stability progressively increased with increasing number of sequential species, becoming the largest in communities composed entirely of sequential species (Fig. 3c, bottom). As before, these results are qualitatively robust to eliminating lag times, but with a weaker effect (Fig. S5). This suggests that for fixed environmental conditions, sequential species would tend to be over-represented diverse niche-packed communities.

## Discussion

In this manuscript, we introduced and measured two different notions of fitness of metabolic strategies in time-varying environments. These notions were (a) the time-averaged growth rate of microbial species using each strategy, and (b) the prevalence of species using each strategy in stochastically assembled microbial communities. Our central result is that sequential strategies become progressively fitter in ecological contexts over co-utilizing strategies using both notions of fitness. The observed fitness difference becomes especially pronounced with increasing community diversity as measured by species richness, as well as the size of the species pool from which communities are assembled.

It is important to understand how the different key parameters in our model affect the strength of this central result. The most explicit of these parameters is the pre-allocation fraction *ϕ*_pre_ which represents the fraction of the proteome that species pre-allocate to resources that they do not consume in a particular environment. The advantage of sequential utilizers persisted for a reasonably broad range of *ϕ*_pre_ (Fig. 2c–e). Specifically, low values of *ϕ*_pre_ increasingly favored sequential species, while higher values favored co-utilizers. The overwhelming advantage of co-utilizers in monoculture contexts comes from their significantly lower lag times. In community contexts, complementary sequential utilizers — with distinct top-choice resources — can assemble to form communities where each species essentially depletes all resources together, and species barely derive any growth from resources other than their top choice. Such collective growth makes the lag disadvantage of sequential species immaterial, contributing to their increased time-averaged growth rate (fitness) in ecological contexts (Fig. 2d).

In addition to *ϕ*_pre_, the lag times depend on the prefactor *τ*_0_ (Methods), which sets an overall scale for lag times. With no lags (*τ*_0_ = 0, we no longer see a stark difference between time-averaged growth rates in monoculture versus ecological contexts (Fig. S5). Sequential species still retain a “biomass advantage” in communities as measured by prevalence (Fig. S5a, green). The advantage in this case can be explained by a rather different effect: namely the statistics of growth rate distributions for species utilizing either strategy (Fig. S5b–c). The odds of a sequential species to win in a head-to-head competition with a co-utilizing species depend on which strategy’s growth rate distribution has the larger median, which at *ϕ*_pre_ ≈0.06 is the co-utilizing strategy (Fig. S5c). However, during community assembly from a large species pool, the level of competition is much greater since only *n*_*R*_ = 4 species can survive out of the pool of *N*_pool_ = 100 introduced species. In this scenario, it is not the medians, but the extreme value of the growth rates that matter. The extremes are determined by the variance of the growth rate distributions, which is larger for the sequential strategy. This is because, unlike co-utilizing strategies, sequential species do not average over *n*_*R*_ growth rates, one for each resource, to determine their growth rate in any environment. Hence, for the same *ϕ*_pre_ ≈ 0.06, it is the sequential strategy, not the co-utilizing strategy, that does better during community assembly (Fig. S5b).

Thus, not just lag times, but the width of the growth rate distributions also contribute to the success of metabolic strategies. In Fig. S3, we systematically investigate the effect of changing this width *σ*, finding that lower values of *σ* lead to a lower fitness of sequential strategies as measured by their prevalence in assembled communities (Fig. S3a). This is consistent with above observation that the success of sequential strategies in communities is determined by the extreme value distributions of growth rates, whose median depends positively on *σ*.

The degree to which the time-varying environments can be considered boom-and-bust environments depends on the dilution factor *D*, with increasing *D* leading to stronger busts. We found that changing *D* had a consistent but small effect of changing the fitness of sequential strategies in community contexts. In particular, lag times become more important for lower values of *D*, since at steady-state, species grow by a smaller overall factor. As a result, as *D* decreases, so does the fitness of sequential utilizers in community contexts as measured by biomass prevalence (Fig. S2a, green).

Our other key result is that communities with sequential species are also significantly more structurally stable, and thus able to withstand greater fluctuations in resource supply ratios. To understand why this is the case, consider the following argument. Sequential species use only one resource at a time. Importantly, some sequential species may never get to utilize an available resource before it gets depleted by others in the community. For example, if the most preferred resource by a species gets depleted last by the community, this species will never get to use another resource, even though it might be capable of using all resources. In contrast, co-utilizing species always consume all available resources regardless of their depletion order. As a consequence, changing the concentration of any resource will affect all co-utilizers but only a fraction of sequential species. In more mathematical terms, the matrix *M* encoding which species consume which resources in the steady state cycle (Supplementary Text) will be sparser for sequential communities than co-utilizing ones and hence be more structurally stable. With increasing number of resources, the *M* matrix would get larger and thus we expect the systematic difference between sequential and co-utilizing species to increase. We explore structural stability more systematically, with its dependence on model parameters, in the supplementary materials (Fig. S2, S3, S4).

### Testable predictions

Our results might potentially explain the diversity of metabolic strategies observed in nature, specifically why sequential (diauxic) and co-utilizing strategies coexist without one systematically out-competing the other. In our model, both strategies stratify in communities with different diversity, or species richness, relative to the number of resources (Fig. 3a,d). This leads to the following testable prediction: niche-packed communities (*n*_*S*_*/n*_*R*_ ≈ ≪ 1) would tend to be richer in sequential species, while poorly packed (*n*_*S*_*/n*_*R*_ ≪ 1) communities would be dominated by co-utilizers. Future experimental or observational work could potentially test this prediction and help explain the diversity of observed metabolic strategies.

Another observation of our work is that the fraction of sequential species in communities increases with the species pool size (Fig. S1). The species pool size represents the “maturity” of a community, i.e., the cumulative number of invasion attempts by other species into a community. We predict that more mature communities in nature will tend to be enriched in sequential species.

## Acknowledgements

We thank Y. Fridman, G. Chure and J. Cremer for valuable discussions. A.G. acknowledges support from the Ashok and Gita Vaish Junior Researcher Award, the Department of Atomic Energy, Government of India, under project number RTI4001, as well as the Government of India’s DBT Ramalingaswami Fellowship.

## Data and code availability

There are no data associated with this paper. All code is available as a GitHub repository at the following link: https://github.com/maslov-group/Ecol_adv_diaux

## Methods

### Model of microbial proteome allocation

To incorporate proteome allocation into our consumer-resource model in boom-and-bust environments, we extended previous proteome allocation schemes to include pre-allocation of metabolic enzymes. That is, just as in previous models, we assumed that the proteome of each individual in a species was composed of metabolic enzymes, ribosomes and other proteins [17]. We perform allocation in two steps. First, we assign the proteome into the ribosomal and metabolic sectors assuming that allocations to other sector(s) are fixed. Second, we subdivide the metabolic sector into distinct enzymes for different resources.

To describe the allocation between metabolic enzymes and ribosomes, we choose to adapt the model of Hermsen et al. [16]. Consider a species *α* growing in a particular resource environment. To grow on these resources, the species allocates a certain fraction of metabolic enzymes 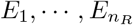, normalized to 1. Note that in our model, species allocate enzymes even towards resources that they are not currently consuming. The growth rate *g*_*α*_ of the species is determined by the following equation:

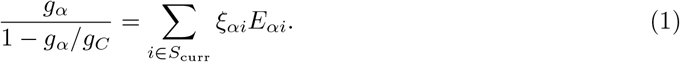

where *S*_curr_ is the set of resources the species is currently consuming. This equation can be rear-ranged to obtain:

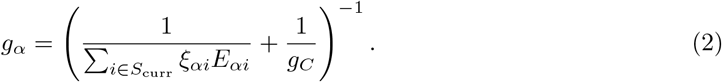

Here, *g*_*C*_ is the theoretical upper bound on the growth rate which only depends on the species. We set it to the same value equal to 1 for all species for simplicity. Further *ξ*_*αi*_ is a species-specific coefficient representing the contribution of given resource *i* towards growth per unit enzyme fraction. In our simulations we select it from a truncated Gaussian distribution with mean 0.5 and standard deviation 0.1.

We then consider how species allocate their proteome towards different metabolic enzymes *E*_*αi*_. This proteome allocation is different for each strategy. Here, we assume that for each resource there is always a small portion of the proteome *ϕ*_pre_ that has been “pre-allocated” to the metabolic enzyme necessary to utilize this resource. This “pre-allocated” fraction of the proteome can never be reallocated to other resources.

For a co-utilizing species with *n* resources present in the environment (*n* ≤ *n*_*R*_), each such resource would initially have *ϕ*_pre_ fraction of the metabolic sector of the proteome pre-allocated to it. Hence, the total fraction of the metabolic enzyme pool that could in principle be reallocated is given by (1− *n*_*R*_*ϕ*_pre_). For co-utilizers, we assume that this amount is distributed among all *n* resources equally (for simplicity). In this case, each of the *n* resources currently utilized by the species has a total fraction of enzymes given by

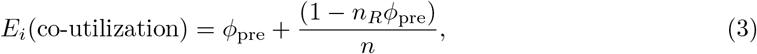

while each resource not currently utilized has a fraction *ϕ*_pre_ allocated to it. For a sequential species *n* = 1 because they always consume one resource at a time. Thus,

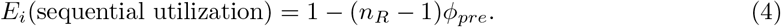

To understand how *ϕ*_pre_ affects the balance between sequential and co-utilizing strategies in head-to-head competition, suppose there are two strains *c* and *s* sharing the same set of physiological parameters (*ξ*_*ci*_ = *ξ*_*si*_ = *ξ*_*i*_), but *c* is a co-utilizing strain, while *s* is a top-smart sequential strain. These strains are grown in a boom-and-bust environment with *n*_*R*_ resources at the beginning of each cycle. When they compete against each other, our previous work shows that the competition occurs mainly during the first temporal niche where all resources are present [4]. To compare the growth rates of the sequential and co-utilizing species before correcting for the ribosomal proteome sector, we only need to compare the RHS in Eq. (1). The ratio *r*_*sc*_ quantifies the advantage of the top-smart sequential strategy over co-utilization one. When all substrates are present it is given by

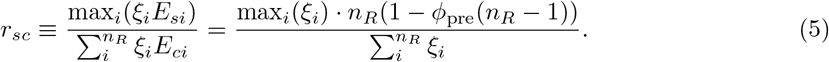

Note that for *ϕ*_pre_ = 0 the sequential strategy always has an advantage in the first temporal niche: *r*_*sc*_ *>* 1. Indeed, in this case *r*_*sc*_ is given by the ratio of the maximum over the mean growth rate, which is always larger than 1. As *ϕ*_pre_ increases, a larger reallocation portion of metabolic enzymes starts to favor the co-utilization strategy over the sequential utilization one, so that at some value of *ϕ*_pre_ the ratio would satisfy *r*_*cs*_ *<* 1. This argument qualitatively explains the behavior of the fraction of surviving sequential utilizers in head-to-head competition shown in Fig. S5a (black curve).

### Model of microbial community dynamics in boom-and-bust environments

We model microbial community dynamics in a boom-and-bust environment where *n*_*R*_ substitutable resources are cyclically supplied at concentrations *R*_*k*_ (*k* = 1, 2, … *n*_*R*_). We attempt to assemble a community by adding species to this environment. Each species is assumed to be a generalist able to grow on each of these resources. We denote species by Greek symbols, so that the abundance of species *α* is represented by *N*_*α*_(*t*); the concentration of resource *k* is represented by *R*_*k*_(*t*). Depending on the metabolic strategy of a species, it consumes certain resources quantified by a time-dependent consumption matrix *c*_*αk*_(*t*). If species *α* consumes resource *k* at time *t*, we set *c*_*αk*_(*t*) to 1; otherwise, we set it to 0.

We assume that resource concentrations are much higher than the species’ half-saturating substrate concentrations during the entirety of all temporal niches in which a resource is present. This is a reasonable assumption as long as initial resource concentrations are much larger than their corresponding half-saturating concentrations. Thus, in our model, in each temporal niche, species grow exponentially at constant growth rates given in the previous section as Eq. (2). Note that *E*_*αk*_(*t*) is a function of *c*_*αk*_(*t*)’s, according to 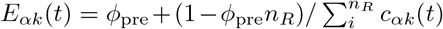,which comes from Eq. (3). Thus, the dynamics of species abundance in our model can be written as follows

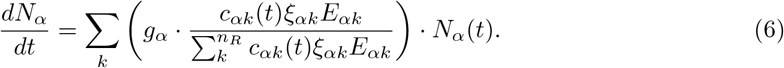

Similarly, the resource depletion dynamics can be written as follows

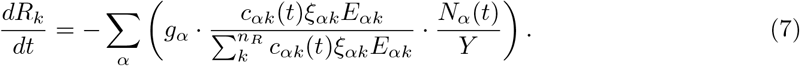

For the sake of simplicity, we define resource concentrations so that the yields *Y* all equal 1.

The “boom” phase of each cycle ends when all resources are depleted. It is then followed by a “bust” phase during which all microbial abundances are reduced (diluted) by the same factor *D >* 1 until the arrival of the next nutrient bolus, when the boom phase resumes. Our dynamical equations are inspired by the protocol of serial dilution experiments performed in many laboratories to study microbial communities in vitro. A version of this dynamics is also realized in natural microbial ecosystems living in boom-and-bust environments characterized by long intervals between bust phases and large boluses delivered at the beginning of each boom phase.

We are interested in computing the steady state of the boom-and-bust dynamics in which each species grows by exactly the same factor *D* by which it is subsequently diluted. To compute this steady state dynamics, it is convenient to divide the boom phase of the cycle into a sequence of *n*_*R*_ *temporal niches* where only a particular subset of resources is present [11, 18, 19]. The total number of possible temporal niches for *n*_*R*_ resources is given by 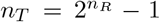.Here we have excluded a temporal niche where all resources are absent and therefore no species can grow. The resources present in the environment disappear one by one, resulting in a particular subset of *n*_*R*_ temporal niches (out of *n*_*T*_ possible ones) realized for each of the *n*_*R*_! orders of resource depletion. For example, if three resources disappear in the order 2 →3 →1, the corresponding temporal niches are 111→ 101→ 100 (again, we do not show or count the last niche 000, which separates the end of the boom phase and the beginning of the bust phase of the cycle).

Note that we could have also assumed that species can die during the temporal niche where all resources are absent, e.g., due to starvation. Indeed, it is easy to generalize our model to incorporate the possibility of species dying (having a negative growth rate) during any temporal niche. In this situation, competition occurs even in the temporal niche where all resources are absent, adding one more temporal niche to *n*_*T*_. Also, there will be *n*_*R*_ + 1 temporal niches between adjacent boom phases.

### Model of lag times

In the version of our model with lag times, we assume that after a resource has been depleted, the species that have been consuming it will enter a lag phase, during which they will reallocate their enzyme pools and will not grow at all. During this phase, the enzymes necessary to utilize a single resource (or multiple resources in the case of co-utilizers) are synthesized by an exponential enzyme synthesis process, starting from its pre-allocated level *ϕ*_*pre*_ and ending when it reaches the maximum amount (1 − (*n*_*R*_ − 1))*ϕ*_*pre*_. This process is based on the assumption that enzymes synthesize precursors (charged tRNA) necessary for the synthesis of additional enzymes of their own kind[15]. We model the process as 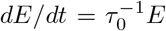,where *E* is the amount of enzyme being synthesized and *τ*_0_ is a time scale. For sequential utilizers, integrating from *t* = 0 to *t* = *τ*, we get a lag time *τ* as

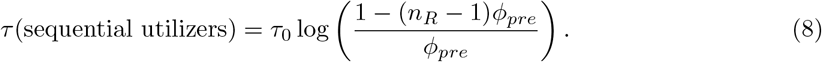

For co-utilizers, the log term will include a ratio of the pre- and post-shift allocation fractions, resulting in much shorter lags compared to sequential utilizers.

In Ref. [5], it was proposed and experimentally confirmed that when species switch from glycolytic to gluconeogenic carbon sources, the dynamics of the enzymes necessary to utilize the latter resources are instead given by 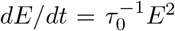; this is due to the peculiar biochemistry of these two complementary pathways. In this case we get a different expression for the lag *τ* of sequential utilizers as

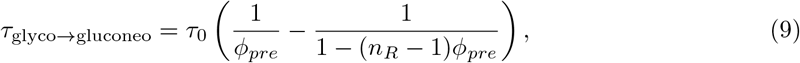

along with a corresponding variant for co-utilizers.

### Details of numerical simulations

In all of our numerical simulations, we sampled enzyme efficiencies from *ξ*_*αi*_ = *ξ*_0_ + *σ*_*ξ*_*Z*_*αi*_, where *Z*_*αi*_ is a random number drawn from the standard normal distribution. For the figures shown in the main text of the study, we used *ξ*_0_ = 0.5 hr^−1^ and *σ*_*ξ*_ = 0.1 hr^−1^. To prevent negative growth rates and to keep the *ξ*_*αi*_ distribution symmetric around its mean, we truncated the normal distribution so that 0 *< ξ*_*αi*_ *<* 1. The upper bound of the growth rate was set to *g*_*C*_ = 1 hr^−1^. All sequential species were set to be top-smart, i.e. with the highest growth rate on their most preferred resource, while their preference order on other resources was randomly generated. The initial lag for each species, where they transition from the dormant state to active growth, was sampled from *τ*_initial_ ∼ U(2, 3) hr. The lags caused by enzyme reallocation were generated from eq.8, where the coefficient *τ*_0_ for each species was sampled from *τ*_0_ ∼ U(0.2, 0.4) hr.

In the simulations described in Fig. 2b, for each value of *ϕ*_pre_, we simulated serial dilution experiments with dilution factor *D* = 1000 and dilution interval *T* = 24 hours. For community assembly, we sampled 200 species pools, each with 50 sequential and 50 co-utilizing species. For head-to-head competition, we sampled 2000 pairs of sequential and co-utilizing species. During community assembly from each species pool, 4 resources added at the beginning of each dilution cycle were sampled uniformly from the simplex Σ_*i*_ *R*_*i*_ = 4 at the beginning and were fixed for each dilution cycle throughout the assembly process. The duration of each dilution cycle is set to 24 h. We attempted to invade the system multiple times with all species from the pool in a random order until no more successful invasions were possible, i.e. a non-invadable state was reached. All invaders from the pool were introduced with a low abundance of 10^−8^, which is much lower than the abundance of any resident species in a steady state community. The elimination bound for a species was also set to 10^−8^.

To generate Figures 2c-d, we sampled 100 sequential and 100 co-utilizing species to measure their time-averaged growth rates under monoculture growth. For full diversity communities, we generated 100 purely sequential and 100 purely co-utilizing communities where *n*_*S*_ = *n*_*R*_ = 4 species coexist (see Supplementary Text). We measured the time-averaged growth rate by ⟨*g*⟩_*T*_ = log*D/T*_dep_, where *T*_dep_ is the time it takes to deplete the last resource during a dilution cycle at steady state. In the structural stability simulations in Fig. 3c, for each possible composition 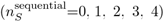, we randomly sampled 10000 communities where *n*_*S*_ = *n*_*R*_ = 4 species coexist (see Supplementary Text). The *ϕ*_*pre*_ for these species was set to 0.036. We then calculated structural stability for these communities (see Supplementary Text).

All simulations were done in Python (see Code Availability Statement).

## Notes

https://github.com/maslov-group/Ecol_adv_diaux

### Competing Interest Statement

The authors have declared no competing interest.

### Summary of Updates

Significant re-analysis and modified framing of the results. New title, abstract, figure 2 and Results.

