## Supplementary Information for "Fitness advantage of sequential metabolic strategies emerges from community interactions in strongly fluctuating environments"

### Supplementary Information
### Emergent ecological advantage of sequential metabolic strategies in complex microbial communities

Zihan Wang[1,2], Yu Fu[3], Akshit Goyal[4,*], Sergei Maslov[1,2,5*],

[1] *Department of Physics, University of Illinois Urbana-Champaign, Urbana, Illinois, USA.*
[2] *Carl R. Woese Institute for Genomic Biology, University of Illinois Urbana-Champaign, Urbana, Illinois, USA.*
[3] *Department of Physics, Yale University, New Haven, Connecticut, 06511, USA*
[4] *International Centre for Theoretical Sciences, Tata Institute of Fundamental Research, Bengaluru 560089, India.*
[5] *Department of Bioengineering, University of Illinois Urbana-Champaign, Urbana, Illinois, USA.*

#### Supplementary Figures

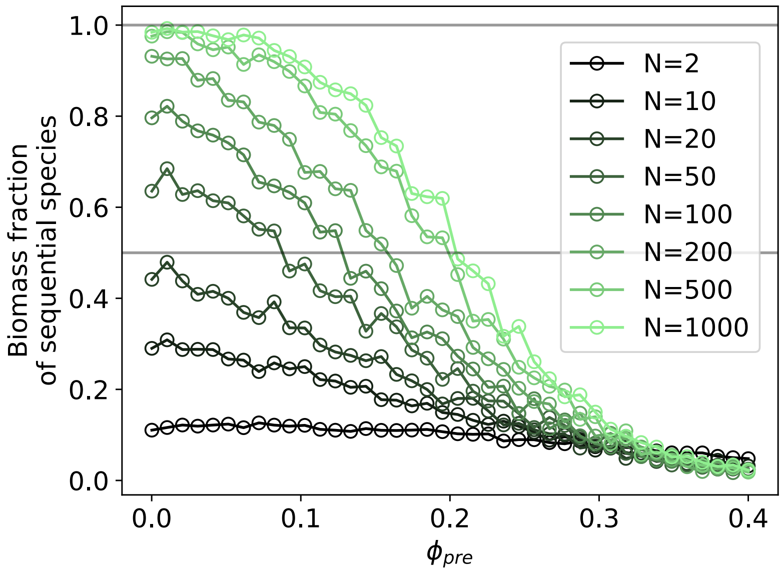


Supplementary Figure 1: **Pool size of assembly affects the ecological advantage of sequential utilizers.** Fraction of sequential utilizers in biomass among survivors (y-axis) are plotted at different pool sizes of the community assembly (represented by different shades of green), as a function of the pre-allocation factor $\phi$. For each value of $\phi$, we randomly generated 200 species pools, and each of them consists of 50% top smart sequential species and 50% co-utilizing species whose growth rates are sampled from the same distribution as in main text (Fig.2d). The gray horizontal line is at $y = 0.5$.

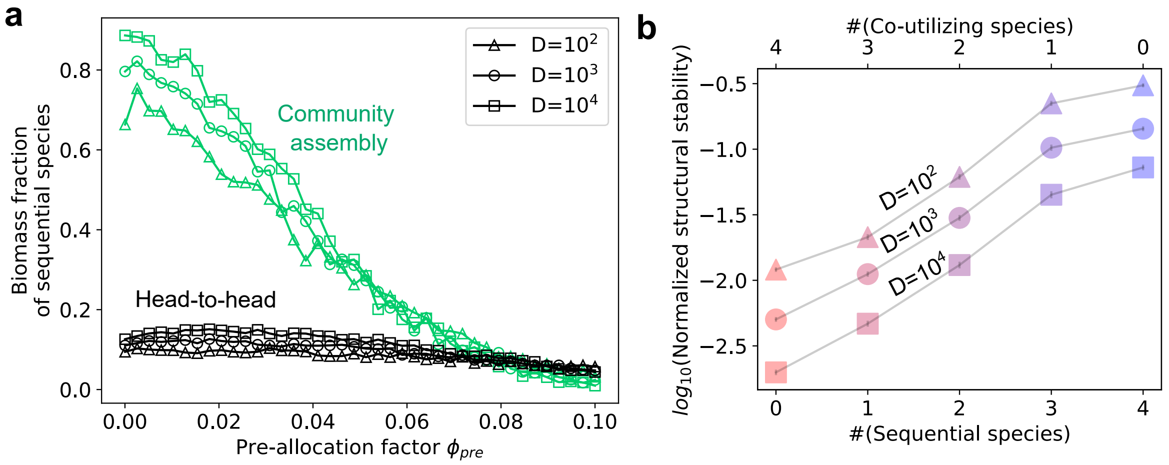


Supplementary Figure 2: **Effect of dilution factor $D$ on the ecological advantage of sequential species.** (a) Fraction of sequential utilizers in biomass among survivors, in each of the two scenarios: head-to-head (black) and in complex communities (green), plotted as a function of the pre-allocation parameter $\phi_{\text{pre}}$ common to all species in the pool. Simulations were performed in the same setup as in Fig. 2e. Marker shapes indicate the dilution factor $D$. The growth period is shorter under lower $D$, making the sequential lag's negative effects more pronounced. (b) The structural stability of niche-packed communities depends on $D$. Simulations were performed in the same way as in Fig. 3c, where the average logarithm of normalized structural stability (y-axis) is plotted as a function of the composition of niche-packed communities ($n_S = n_R = 4$). Marker shapes represent the value of $D$, and colors reflect the number of sequential/co-utilizing species. When generating these communities $\phi_{\text{pre}}$ was taken at 0.036 (Fig. 3b).

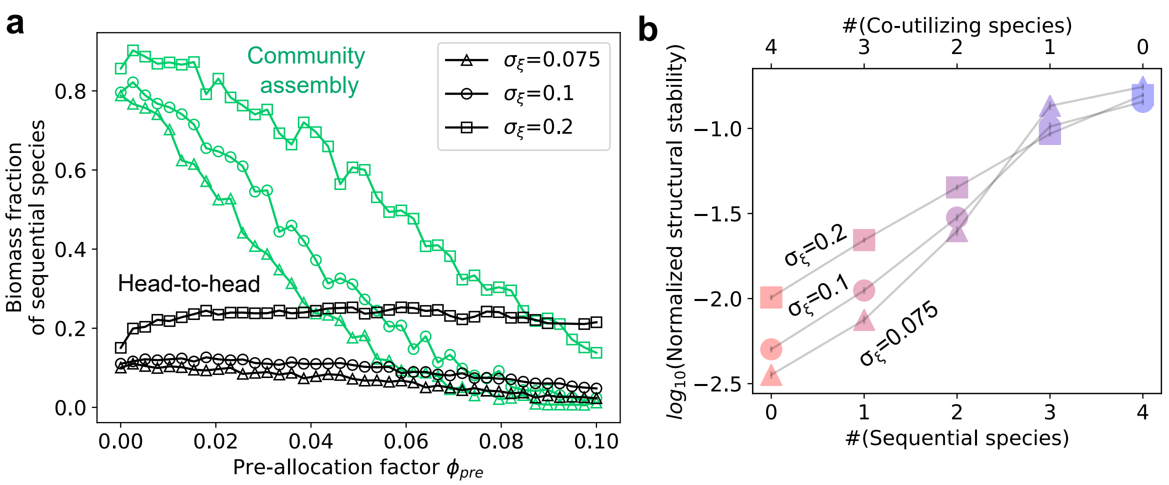


Supplementary Figure 3: **Effects of growth rate distribution on the ecological advantage of sequential species.** (a) Fraction of sequential utilizers in biomass among survivors, in each of the two scenarios: head-to-head (black) and in complex communities (green), plotted as a function of the pre-allocation parameter $\phi_{\text{pre}}$ common to all species in the pool. Simulations were performed in the same setup as in Fig. 2e. Marker shapes indicate the width of enzyme efficiency distribution $\sigma_\xi$ of $\xi_{\alpha i}$, which contributes to the growth rates. (b) The structural stability of niche-packed communities depends on $\sigma_\xi$. Simulations were performed in the same way as in Fig. 3c, where the average logarithm of normalized structural stability (y-axis) is plotted as a function of the composition of niche-packed communities ($n_S = n_R = 4$). Marker shapes represent the value of $\sigma_\xi$, and colors reflect the number of sequential/co-utilizing species. When generating these communities $\phi_{\text{pre}}$ was taken at 0.036 (Fig. 3b).

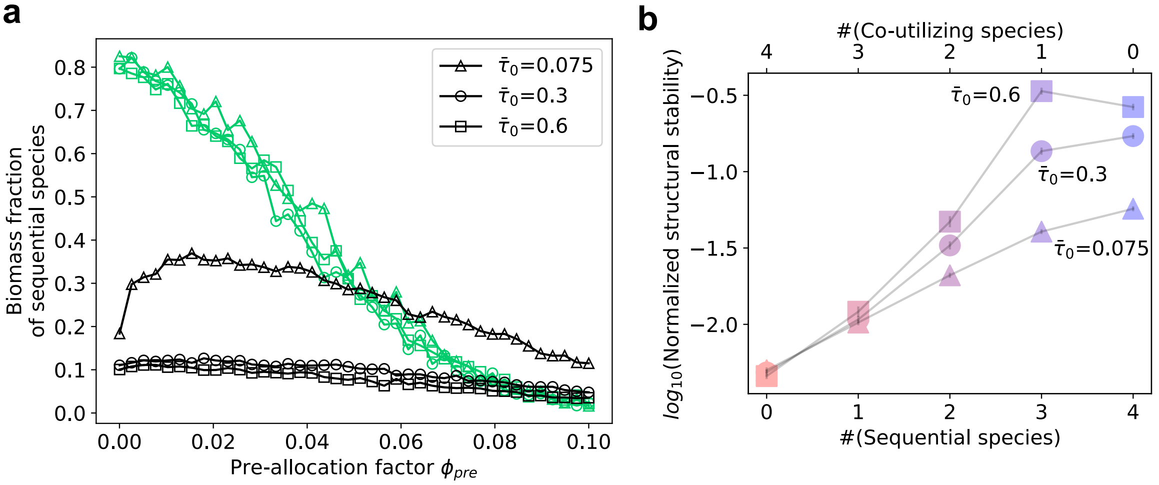


Supplementary Figure 4: **Effects of lags on the ecological advantage of sequential species.** (a) Fraction of sequential utilizers in biomass among survivors with different lags: head-to-head (black) and in complex communities (green), plotted as a function of the pre-allocation parameter $\phi_{\text{pre}}$ common to all species in the pool. Simulations were performed in the same setup as in Fig. 2e. Marker shapes indicate the mean coefficient $\bar{\tau}_0$ of proteome reallocation lag. The sequential lag is calculated by $\tau = \tau_0 \log\left(\frac{1-(n_R-1)\phi_{pre}}{\phi_{pre}}\right)$ where the coefficient $\tau_0$ is sampled from a uniform distribution $\bar{\tau}_0 \cdot \text{U}(0.67, 1.33)$ (Methods). (b) The structural stability of niche-packed communities depends on $\bar{\tau}_0$. Simulations were performed in the same way as in Fig. 3c, where the average logarithm of normalized structural stability (y-axis) is plotted as a function of the composition of niche-packed communities ($n_S = n_R = 4$). Marker shapes represent the value of $\bar{\tau}_0$, and colors reflect the number of sequential/co-utilizing species. When generating these communities $\phi_{\text{pre}}$ was taken at 0.036 (Fig. 3b).

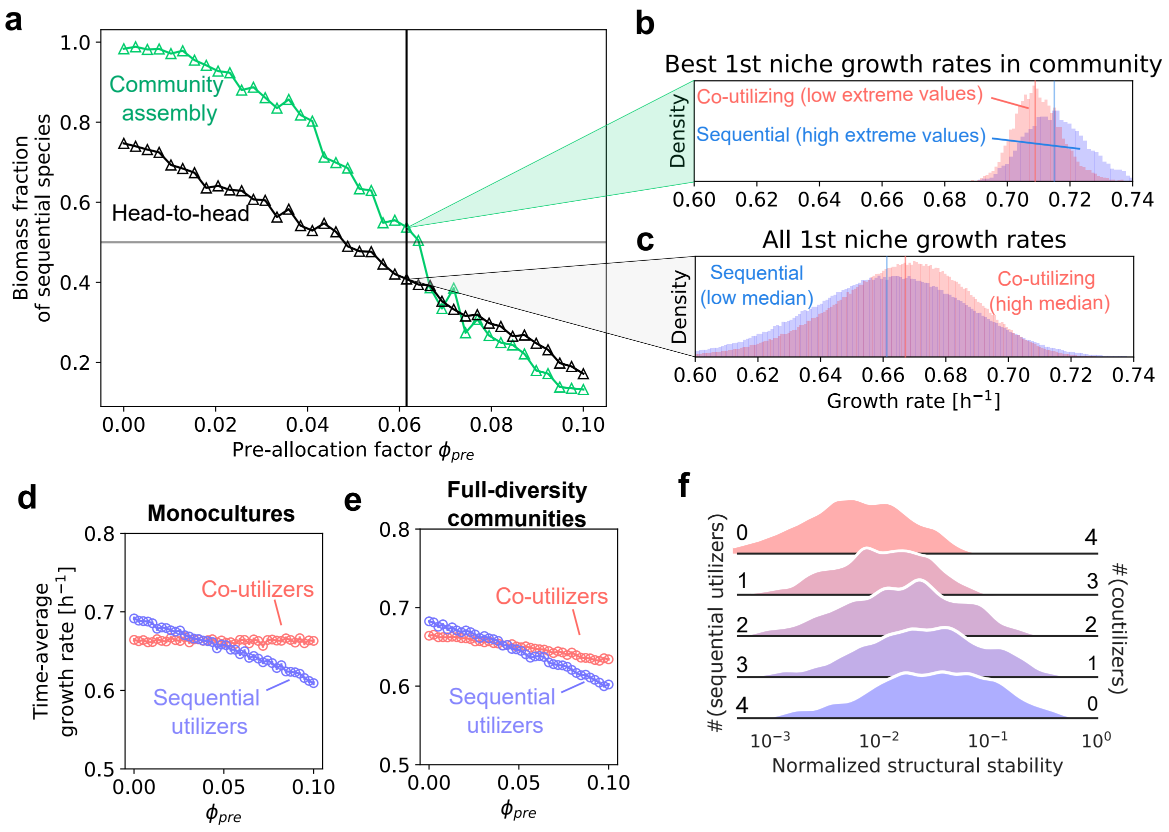


Supplementary Figure 5: **Ecological effects of metabolic strategies when lags are absent.** (a) Fraction of surviving sequential utilizers in each of the two scenarios: head-to-head (black) and in complex communities (green), plotted as a function of the pre-allocation fraction $\phi_{\text{pre}}$ common to all species in the pool. Lags strongly affect the pairwise competition outcomes of sequential species: compared to Fig.2f, their biomass fraction increases a lot, while in assembled communities it does not increase as much. The gray horizontal line is at $y = 0.5$. The black vertical line lies at $\phi_{\text{pre}} = 0.0615$, where sequential species hold a slight advantage in communities, but a slight disadvantage in head-to-head competitions. (c) Distribution of growth rates of sequential (blue) and co-utilizing (red) species in the first temporal niche for $\phi_{\text{pre}} = 0.0615$. Sequential species have a lower mean and higher variance. Vertical lines are medians of the corresponding distributions. (b) Distribution of top growth rates selected from each of the distributions in (c); sequential species have greater extreme values due to wider upper tails and thus win in complex communities where competition is strong. Vertical lines are medians of the corresponding distributions.(d)-(e) Time-average growth rates of communities as a function of $\phi_{\text{pre}}$. (d) In monocultures, the sequential utilizers (blue) gain a growth advantage over coutilizers (red) at low $\phi_{\text{pre}}$, which is also reflected in (a). (e) In fully-packed communities, the trend is similar. (f) Distributions of normalized structural stability for niche-packed communities ($n_S = n_R$ species). Apart from the lag time being zero, all other parameters used to generate these communities are identical to the ones in Fig. 3c. Increasing the number of sequential utilizers from 0 to 4 systematically increases structural stability.

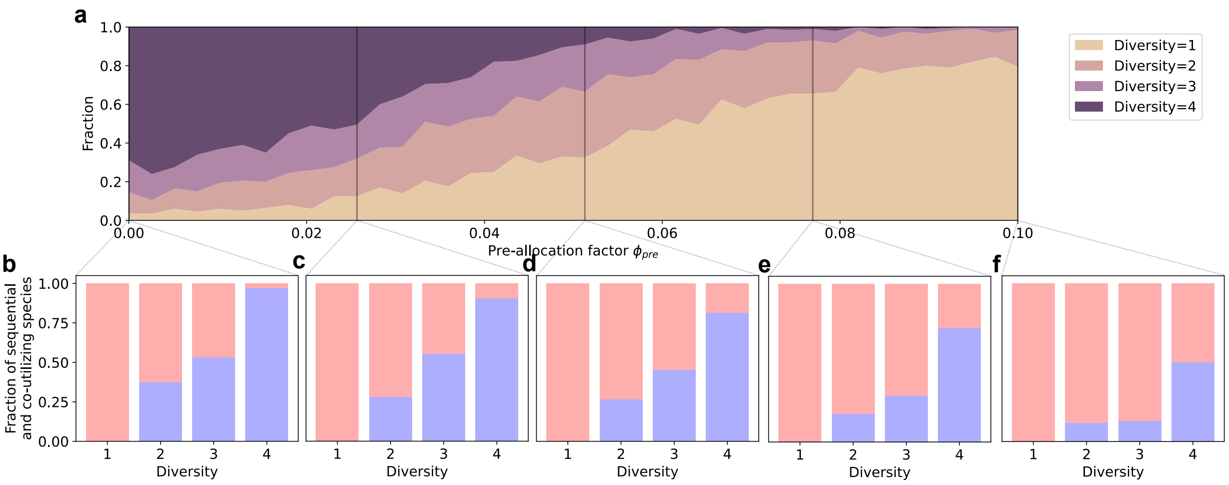


Supplementary Figure 6: **Composition of assembled communities stratified by diversity across different values of $\phi_{\mathbf{pre}}$.** (a) Same as Fig. 3b. (b)-(f) Stacked bar plot showing how sequential (red) and co-utilizing species (blue) are stratified across communities with different diversities, at specific $\phi_{\text{pre}}$ values marked on (a).

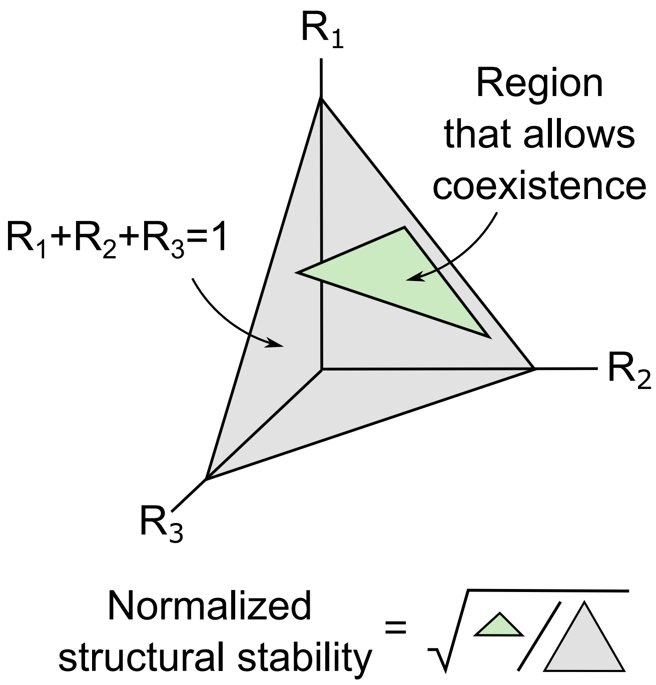


Supplementary Figure 7: **Schematic of structural stability.** Structural stability quantifies the fraction of supplied resource concentration ratios in which a feasible community can assemble (see Supplementary Text). It is defined as the fraction of resource supply ratios that support the coexistence of a given microbial community (green) in all possible resource supply ratios (gray).

#### Supplementary Text

##### The feasibility of a community assembly

The steady state of a microbial community in a boom-and-bust environment is characterized by a particular depletion order of resources which depends on abundances of both species and resources at the start of the boom cycle in a complex fashion. Since it is not known a priori in which order the resources might be depleted, one should test all $n_R!$ possible depletion orders for feasibility. For a given resource depletion order let $G_{\alpha i}$ be the growth rate of the species $\alpha$ in the temporal niche $i$ ranging from $i = 1$ where all $n_R$ resources are present to $i = n_R$ where only one resource remains. Let $t_i$ be the duration of each of these temporal niches. In the absence of time lags we can calculate the growth ratio of each species during the entire boom phase of the cycle as $\exp(\sum_{i=1}^{n_R} G_{\alpha i} t_i)$. In the steady state, this ratio must be equal to the dilution factor $D$ during the bust phase of the cycle, which leads to the following equations for each of $n_S$ species

$$\sum_{i=1}^{n_R} G_{\alpha i} t_i = \log D. \tag{S1}$$

The matrix $G_{\alpha i}$ used in this equation is related to the matrix $g_{\alpha k}$ of growth rates on individual resources, and this relationship depends on the metabolic strategy of the species. If the matrix $G_{\alpha i}$ is invertible, the above equation can be solved, and the solution is biologically and physically feasible provided that all time $t_i > 0$.

In the presence of lags, the steady-state equation can be rewritten as

$$\sum_{i=1}^{n_R} G_{\alpha i} \cdot (t_i - \tilde{\tau}_{\alpha i}) \cdot \Theta(t_i - \tilde{\tau}_{\alpha i}) = \log D, \tag{S2}$$

where $\Theta(x)$ is the Heaviside step function, and $\tilde{\tau}_{\alpha i}$ is the modified lag time of species $\alpha$ during temporal niche $i$. In most scenarios, $\tilde{\tau}_{\alpha i} = \tau_{\alpha i}$, where $\tau_{\alpha i}$ is the lag time directly derived from the availability of resources during the temporal niche and the metabolic strategy of the species.

A scenario where $\tilde{\tau}_{\alpha i} \neq \tau_{\alpha i}$ can arise as follows: consider an environment where resources are depleted sequentially in the order $1 \to 2 \to 3$. Suppose there is a species $\alpha$ with a resource preference order of $1 \to 3 \to 2$. Under short lag phases, species $\alpha$ would switch from resource 1 to resource 3 during the second temporal niche ($t_2$) and would not switch during the third niche ($t_3$). However, if the lag phase is so long that species $\alpha$ remains in the lag phase throughout $t_2$ ($\tau_{\alpha 2} > t_2$) and only switches during $t_3$, we must adjust $\tilde{\tau}_{\alpha 3}$ accordingly. Specifically, $\tilde{\tau}_{\alpha 3}$ is corrected as $\tilde{\tau}_{\alpha 3} = \tau_{\alpha 2} - t_2$.

In contrast, for a species $\beta$ with a preference order of $1 \to 2 \to 3$, the lag time during $t_2$ does not contribute to reducing its lag phase in $t_3$, because in $t_3$ it switches to a new resource than in $t_2$. Even if $\tau_{\beta 2} > t_2$, species $\beta$ must undergo a full switching process to transition to resource 3 during $t_3$. Thus, for species $\beta$, the corrected lag time remains $\tilde{\tau}_{\beta 3} = \tau_{\beta 3}$.

With the corrections of $\tilde{\tau}_{\alpha i}$ described above, we used an iterative algorithm to solve for the temporal niches $t_i$ (see code for details). In principle, after performing this algorithm across all $n_R!$ depletion orders for feasibility, one can get more than one feasible solution, corresponding to different ways of assembling a niche-packed microbial community with the same set of species.

##### Structural stability with respect to resources

A feasible community with full diversity, given the depletion order of resources, has a unique set of temporal niche durations $t_i > 0$. The next step is to find the range of relative resource concentrations $R_i$ in the nutrient bolus that lead to community assembly. This can be done using the mass conservation rules in the resource-to-biomass conversion. The temporal niche time intervals $t_i$ and lag times $\tilde{\tau}_{\alpha i}$ fully determine the factors $F_{\alpha i} = \exp\left[G_{\alpha i} \cdot \Theta(t_i - \tilde{\tau}_{\alpha i}) \cdot (t_i - \tilde{\tau}_{\alpha i})\right]$, by which individual microbial species grow during that temporal niche. During this time, the biomass of the species $\alpha$ increased from $N_\alpha(0) \prod_{j<i} F_{\alpha j}$ to $N_\alpha(0) F_{\alpha i} \prod_{j<i} F_{\alpha j}$. This increase in biomass is equal to the total resources consumed during this temporal niche. The amount of resource $R_k$ consumed is given by $N_\alpha(0) c_{\alpha k}(\text{during niche i})(F_{\alpha i} - 1) \prod_{j<i} F_{\alpha j}$. This allows one to compute the $M_{\alpha k}$ matrix, which converts species abundances $N_\alpha(0)$ at the start of the boom phase into resource quantities they consumed during the entire boom phase.

$$M_{\alpha k} = \sum_{i=1}^{n_R} \frac{c_{\alpha k}(\text{during niche i}) g_{\alpha k}}{\sum_{k'=1}^{n_R} c_{\alpha k'}(\text{during niche i}) g_{\alpha k'}} (F_{\alpha i} - 1) \prod_{j<i} F_{\alpha j}. \tag{S3}$$

Due to mass conservation column sums are given by $\sum_{k=1}^{n_R} M_{\alpha k} = D - 1$. Indeed, in the steady state, the abundance of each species must increase by a factor $D$. This extra biomass given by $N_\alpha(0)(D-1)$ must be equal to the total quantity $N_\alpha(0) \sum_{k=1}^{n_R} M_{\alpha k}$ of all resources consumed by this species.

The structural stability $S_R$ of a feasible community can be quantified by the fraction of all bolus resource ratios for which the community successfully assembles. It is proportional to the determinant $|\det(M)|$. To properly normalize it, one must first divide the matrix $M$ by $D-1$ so that the sum of the elements in each column is equal to 1. If one randomly chooses $n_R - 1$ ratios between resource concentrations $R_k$ in the nutrient bolus at the beginning of each boom phase of the cycle, the fraction of the volume of the simplex $\sum_k R_k = 1$, $R_k > 0$ that results in community assembly is given by

$$S_R = \frac{|\det(M)|}{(D-1)^{n_R}} \tag{S4}$$

The structural stability defined in this way exponentially scales with the number $n_R - 1$ of independent ratios between resource concentrations. The normalized stability defined by

$$s_R = (S_R)^{1/(n_R - 1)} \tag{S5}$$

corrects for this effect (see Fig. S7). As can be seen from this study, for $D = 100$, $g_0 = 1$ and $\sigma_g = 0.2$ used in our study, the average value of $s_{\text{str R}}$) in communities composed of sequentially utilizing species (both smart and random) is approximately independent of $n_R$. The intuitive interpretation of this quantity is the approximate range of individual nutrient ratios that result in successful community assembly.

In a general case normalized structural stability depends on $n_R$ (the number of resources in the environment), the dilution ratio $D$, the average growth rate $g_0$, and the distribution of growth rates in the $g_{\alpha i}$ matrix.
